# Targeting the glucose-insulin link in head and neck squamous cell carcinoma induces cytotoxic oxidative stress and inhibits cancer growth

**DOI:** 10.1101/2023.07.13.548944

**Authors:** Simbarashe Mazambani, Joshua H. Choe, Tae-Gyu Oh, Pankaj K. Singh, Jung-whan Kim, Tae Hoon Kim

**Affiliations:** Department of Biological Sciences, The University of Texas at Dallas, Richardson TX 75080, USA; Division of Medical Sciences and Department of Medical Oncology, Dana-Farber Cancer Institute and Harvard Medical School, Boston, MA 02115 USA; Department of Oncology Science, The University of Oklahoma College of Medicine, Stephenson Cancer Center, Oklahoma City, OK 73104 USA

**Author notes:** **Corresponding authors:** Jung-whan Kim, Department of Oncology Science, The University of Oklahoma College of Medicine, Stephenson Cancer Center, Oklahoma City, OK 73104 USA., Tae-Hoon Kim, Department of Biological Sciences, The University of Texas at Dallas, Richardson TX 75080, USA.

## Abstract

Reactive oxygen species (ROS) are a double-edge sword in cancers and can both promote pro-tumorigenic signaling and also trigger oxidative stress dependent cell death. Thus, maintaining redox homeostasis to control levels of ROS within a tumor-promoting range elicits critical tumorigenic potential in cancer. Here, we show that head and neck squamous cell carcinoma (HNSCC) is uniquely characterized by its critical dependence on heightened antioxidant capacity facilitated by elevated glucose uptake to maintain survival and proliferation. Using a basal-epithelial-layer-specific GLUT1 knockout mouse model, we establish that targeting GLUT1-mediated glucose utilization in HNSCC cells of origin robustly inhibits HNSCC progression, providing strong genetic evidence that GLUT1 is indeed a targetable metabolic vulnerability. We further demonstrate that disrupting redox homeostasis with prooxidants such as high dose vitamin C and Auranofin induces potent cytotoxicity in HNSCCs by exerting profound oxidative stress when combined with GLUT1 inhibitors. Given the central role of insulin signaling in glucose homeostasis, we additionally show that circulating insulin levels modulate metabolic and oncogenic pathways of HNSCCs, providing a new perspective on events driving and sustaining HNSCC malignancy. These results establish GLUT1 as a viable therapeutic target for HNSCC in combination with prooxidant chemotherapies and define critical dependencies in HNSCC that can be utilized with existing clinical stage drugs for the treatment of HNSCC and potentially other squamous cancers.

## INTRODUCTION

Head and neck squamous cell carcinoma (HNSCC) is one of the most common solid cancers and the sixth leading cause of cancer mortality worldwide, with a reported 890,000 new cases and 450,000 deaths in 2018 (1). HNSCCs arise from the mucosal epithelium of the oral cavity, pharyngeal regions (nasopharynx, oropharynx, hypopharynx) and larynx (2). Epidemiological studies report a diverse range of risk factors for HNSCC including tobacco consumption, alcohol consumption, exposure to environmental pollutants and infection with viral agents, such as HPV and EBV (1,2). To date, no screening strategy has been proven to be effective, and careful physical examination remains the primary approach for early detection. Currently, surgery combined with adjuvant radiation or chemotherapy plus PD-1/PD-L1 antagonists or radiation therapy remain the standard of care for both early and advanced HNSCC (1,3–7). Nevertheless, HNSCC patients have had unsatisfactory outcomes from available therapeutic options.

Our recent work showed an exquisite dependence of squamous cell carcinomas (SCCs) on glucose, driven by highly up-regulated glucose transporter 1 (GLUT1) expression (8). We demonstrated that GLUT1-mediated glucose reliance is a defining phenotype and a unifying feature of all SCCs and established enhanced glycolytic metabolism as an integral process for growth, nourishment and survival of SCCs. Increased glucose influx into squamous cancer cells enhances their antioxidative capacity via the pentose phosphate pathway (PPP) and *de novo* serine biosynthesis (9). Although constitutively elevated oxidative stress is often a key initiator of cancerous lesions, there have been several contradicting studies on the benefits of antioxidants as cancer therapies due to their profound adverse effects (10,11). Various studies have also debated the efficacy of prooxidants as cancer therapies (12–16), however data on the overall benefit of these therapies in HNSCC patients remains largely inconclusive. We sought to understand whether shifting redox homeostasis towards enhanced oxidative stress could lead to significant cell death and attenuate HNSCC growth. Given the delicate balance between cancer initiation and oxidative stress-induced cell death in HNSCCs, we reasoned that HNSCCs might be susceptible to uncontrolled levels of oxidative stress via combinatorial inhibition of glucose influx and treatment with therapeutic prooxidants (17).

The connection between glucose metabolism, redox homeostasis, and insulin signaling has been intensively investigated. Insulin is an anabolic hormone that effectively decreases blood glucose levels by both promoting glucose uptake and utilization in skeletal muscle and adipose tissues, where glycogenesis and lipogenesis then occur, respectively, and preventing gluconeogenesis in hepatocytes. In order to regulate these processes, insulin activates signaling pathways involved in glycolytic metabolism, energy storage, and cell growth. Upon activation, the insulin and insulin-like growth factor (IGF) receptors activate the phosphatidylinositol 3ʹ-kinase (PI3K) and mitogen-activated protein kinase (MAPK) pathways, which regulate cell proliferation, survival, migration, metabolism, and angiogenesis (18–20). Given its vast regulatory impact, insulin signaling plays significant tumor-promoting roles in most cancer types and both hyperinsulinemia and hyperglycemia have been linked to tumor development (21,22). In HNSCCs, *PIK3CA*, which encodes the catalytic subunit of PI3K, is frequently amplified (∼30%) (23) and the resultant highly active PI3K pathway drives glucose uptake by promoting GLUT1 translocation to the plasma membrane among other pro-proliferative cellular outcomes (23–25). Our previous data showed that indirectly suppressing insulin signaling by restricting glucose uptake both perturbed glucose metabolism and suppressed PI3K/AKT signaling in SCC tumors, resulting in decreased SCC tumor growth (9,26). These significant findings motivated our efforts to further probe this glucose-insulin intersection with the goal of yielding new insight into SCCs in diabetic patients, and to understand if direct modulation of circulating insulin levels may provide therapeutic benefits for HNSCC patients. Our current study reveals a tightly coordinated relationship between oncogenic insulin signaling and increased glucose metabolism, which together drive the rapid influx of glucose to fuel the antioxidative demands of squamous cancers.

## RESULTS

### GLUT1 overexpression in TCGA NHSCC tumor tissues

We have previously shown that HNSCC is the highest GLUT1 expressing cancer among over 33 different human cancer types annotated in TCGA database (9). We further analyzed TCGA RNA-seq data from HNSCC patients and matched normal tissue samples. Analyses indicated robust upregulation of GLUT1 (*SLC2A1*), a key rate-limiting factor in the transport of glucose in cancer cells, and genes involved in central glucose metabolism (*HK2*, *ENO1*, *PKM*, *LDHA*), pentose phosphate pathway (*G6PD*), and *de novo* serine biosynthesis pathway (*PSAT1*, *PSPH*, *SHMT2*) (Figure 1). These TCGA analyses indicate that GLUT1 is overexpressed along with genes involved in redox homeostasis in HNSCC.

**Figure 1.**
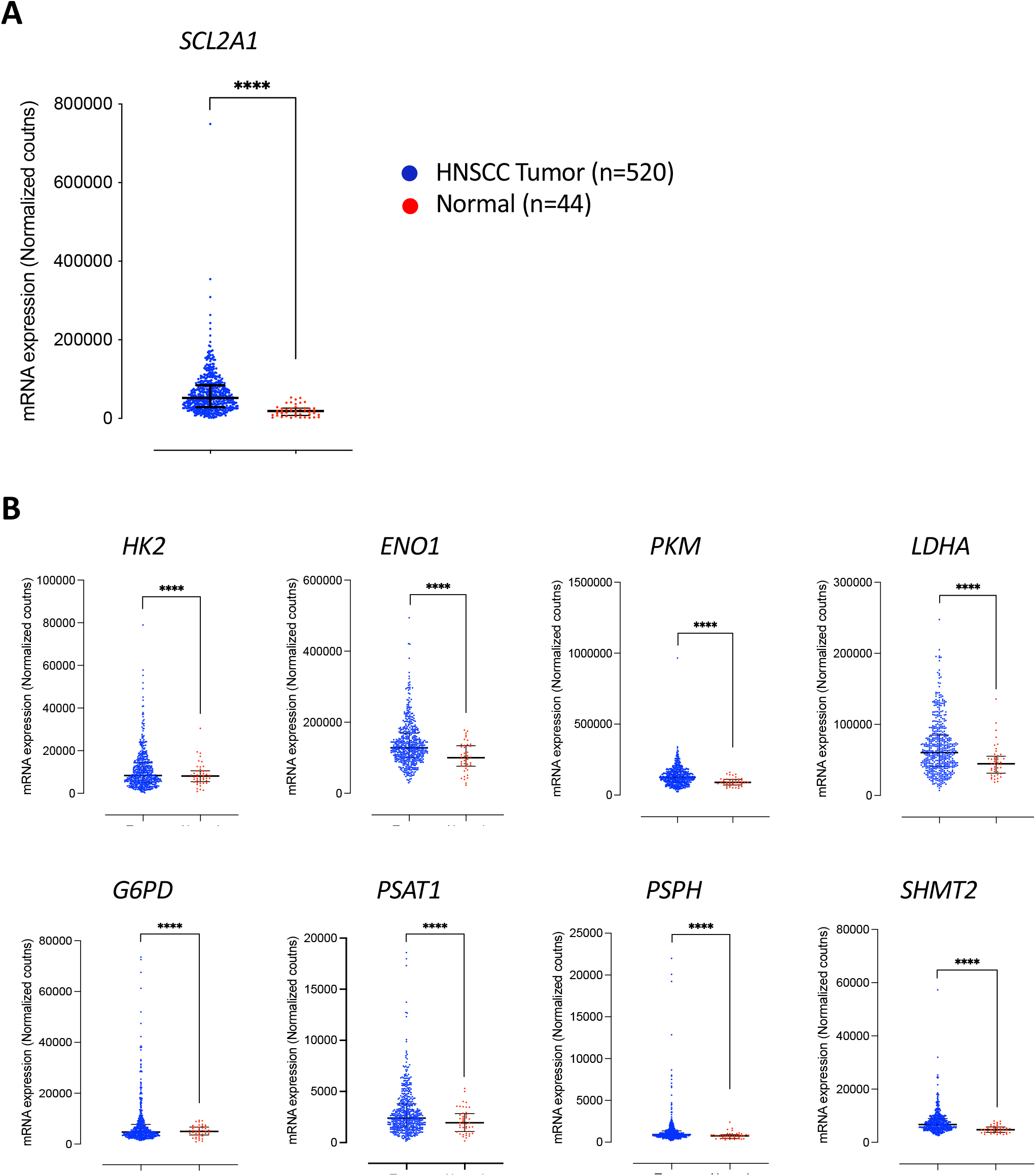
GLUT1 overexpression in TCGA NHSCC tumor samples. TCGA mRNA-sequencing analyses of GLUT1 (A) and metabolic genes associated with glycolysis, pentose phosphate pathway, and *de novo* serine biosynthesis pathway (B) in the TCGA cohort of HNSCC patients (n=520) and matched normal (n=44) tissue samples. Error bars represent the median ± quartile range, Mann-Whitney *u*-test. \*\*\*\**P*<0.0001. SLC2A1, solute carrier family 2 member 1; HK2, hexokinase 2; ENO1, enolase 1; PKM, pyruvate kinase M; G6PD, glucose-6-phosphate dehydrogenase; PSAT1, phosphoserine aminotransferase 1; PSPH, phosphoserine phosphatase; SHMT2, serine hydroxymethyltransferase 2.

### GLUT1 KO attenuates tumor growth in 4NQO mouse model of HNSCC

We next sought to assess whether GLUT1 overexpression in human HNSCC may be recapitulated in the well-characterized 4-nitroquinoline-1-oxide (4NQO)-induced animal model of human oral SCC, a subset of HNSCC. 4NQO is a synthetic water-soluble compound that has been shown to cause DNA adduct formation and induce intracellular oxidative stress thereby mimicking chronic tobacco smoke inhalation and excessive alcohol intake, which are major etiological contributors to oncogenicity in HNSCC (27–29). Histological analysis of tongue tissue harvested from mice exposed to 4NQO in the drinking water (100 ug/mL) for 8 weeks revealed a sequential progression to SCC, starting from hyperplasia followed by dysplasia and invasive SCC (Figure 2A – 2B). Immunohistochemical analysis confirmed the expression of squamous lineage markers, keratin 5 (Krt5, CK5) and p63 in 4NQO-induced tumors (Figure 2B). Tongues collected from mice treated with 4NQO for 4 weeks exhibited hyperkeratosis in the mucosal epithelium, which progressed to moderate-to-severe hyperplasia observed in mouse tongues collected 6 – 10 weeks after exposure to 4NQO (Figure 2B). Severe dysplastic lesions and incidences of invasive SCC were observed in mouse tongues collected 12 – 16 weeks after initial exposure to 4NQO (Figure 2B – 2D). Notably, during the development of premalignant lesions, we observed a dramatic increase in GLUT1 expression in basal epithelial cells, suggesting that GLUT1 upregulation may be an early event during progression to malignancy (Figure 2B).

**Figure 2.**
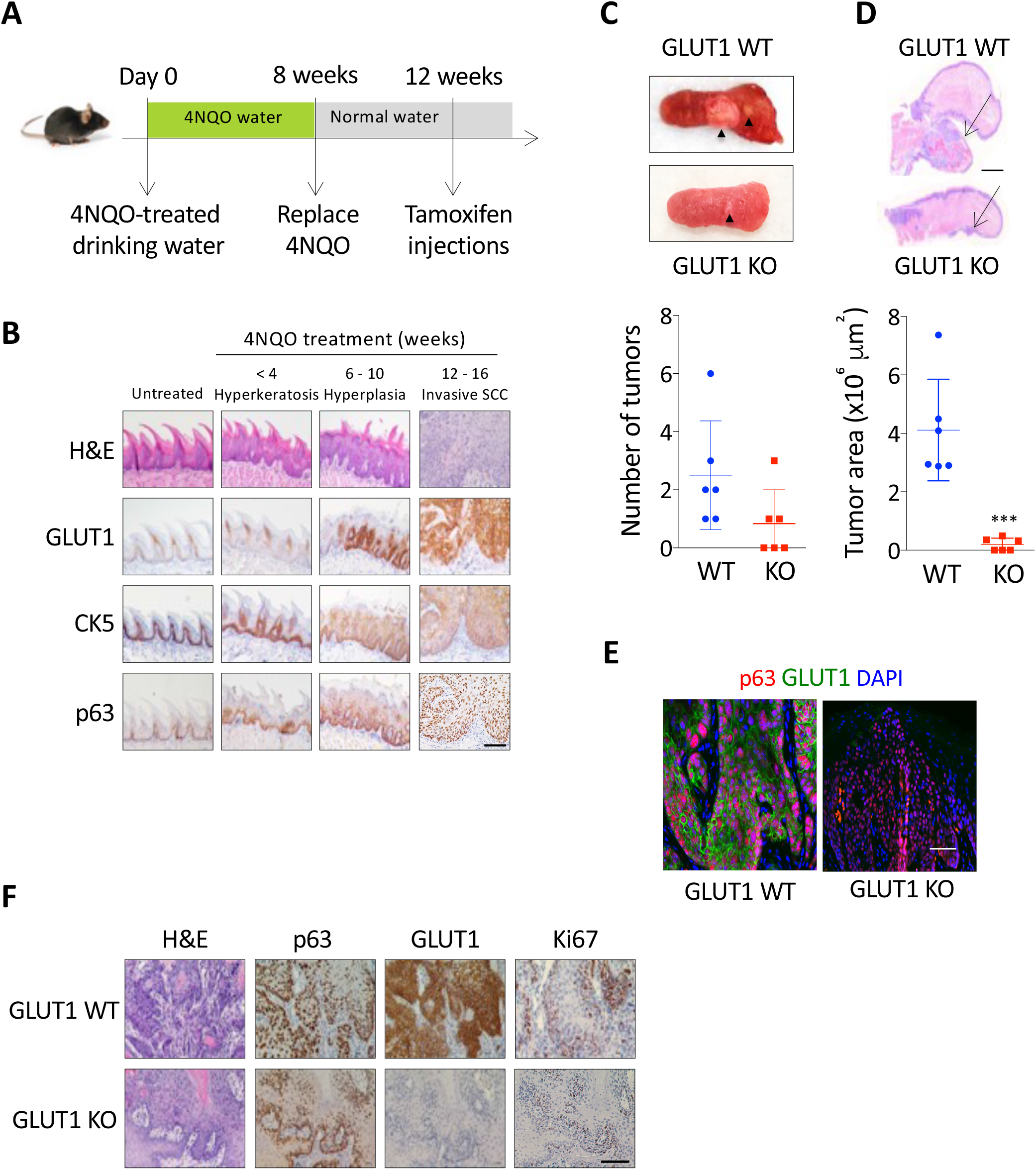
GLUT1 KO attenuates tumor growth in 4NQO mouse model of HNSCC. (A) Schematic showing timeline for treatment of mice with 4NQO. 4NQO was administered in drinking water for 8 weeks and replaced with normal drinking water for the remainder of the study. Tongues were collected every two weeks after first exposure to 4NQO for analyses. For Cre-mediated GLUT1 ablation in basal epithelial cells, tamoxifen was injected i.p. (100 mg/Kg/day for 5 consecutive days) 4 weeks after final administration of 4NQO. Tumors were harvested 4 weeks after the final tamoxifen injection. (B) Representative immunohistochemical analyses of GLUT1 expression in CK5^+^ and p63^+^ basal epithelial cells of untreated and 4NQO-treated mice. (C) Representative images of tumor-bearing tongues from 4NQO-treated mice; black arrowheads depict squamous tongue tumors (Top). Graph shows quantification of number of visible tongue tumors identified in 4NQO-treated mice (n=6) (Bottom). (D) Representative H&E images of 4NQO-treated mouse tongues. Arrows depict 4NQO-induced squamous tumors, scale bar = 1 mm (Top). Graph shows quantification of 4NQO-induced tongue tumor area (μm^2^) in GLUT1 WT and KO mice (n=6), *t*-test (Bottom). (E) Representative co-immunofluorescent staining of GLUT1 and p63 in 4NQO-induced squamous tongue tumors of GLUT1 WT and GLUT1 KO mice. (F) Representative immunohistochemical analyses of p63, GLUT1 and Ki67 expression in 4NQO-induced squamous tongue tumors. \*\*\**P*<0.001. All scale bars = 1 mm.

To investigate whether genetic ablation of GLUT1 in the Krt5+ basal progenitor cells, potential cells of origin of SCC, might effectively inhibit or halt squamous cancer progression post tumor initiation, we established a tamoxifen-inducible knockout mouse model *Krt5:creER; LSL-dTomato; TP53^+/-^; GLUT1^loxP/loxP^* (detailed in methods). This model allowed us to evaluate the impact of basal epithelial cell-specific GLUT1 ablation in HNSCC. *Krt5:creER; LSL-dTomato; TP53^+/-^; GLUT1^loxP/loxP^* (GLUT1 KO) and *Krt5:creER; LSL-dTomato; TP53^+/-^; GLUT1^+/+^* (GLUT1 WT) mice were administered 4NQO-treated drinking water for 8 weeks to induce SCC tumors (Figure 2A). At 16 weeks after initial exposure to 4NQO, all GLUT1 WT mice developed visible tumors on the dorsal or ventral surface of the tongue (arrowheads in Figure 1C depict two visible tumors), whereas, 50% of the GLUT1 KO mice showed no visible tumors, and the remaining GLUT1 KO mice had significantly smaller tumors relative to the GLUT1 WT mice (Figure 2C – 2D). Co-immunofluorescent staining as well as immunohistochemical analysis on serial sections confirmed the specific deletion of GLUT1 in p63^+^ squamous cancer cells in the GLUT1 KO mice (Figure 2E). Furthermore, GLUT1 KO tumor cells exhibited markedly decreased ki67 nuclear staining, indicating a critical role for GLUT1 in SCC cell proliferation (Figure 2F). Collectively, these results suggest that GLUT1 overexpression in the basal progenitor cells is a crucial, and early step necessary for malignant squamous cancer progression and proliferation.

### GLUT1 Inhibition Induces Cytotoxic Oxidative Stress in HNSCC Cells

To elucidate the functional role of GLUT1 in human HNSCC, we employed both genetic and pharmacological approaches to inhibit GLUT1 activity. Knockdown of GLUT1 via shRNA in the FaDu human HNSCC cell line resulted in dramatically impaired cell proliferation (Figure 3A). Additionally, GLUT1 knockdown led to a significant reduction in glucose uptake, which was accompanied by an increase in oxidative stress in the HNSCC FaDu cells (Figure 3B – 3C). These results align with our previous reports and indicate that HNSCC cells heavily rely on GLUT1-mediated glucose uptake to sustain their antioxidative power (8,9). To confirm that the cellular antioxidative capacity driven by glucose intake contributes to the proliferation and survival of HNSCC cells, we treated GLUT1-deficient HNSCC cell line SCC61 with the antioxidant N-acetylcysteine (NAC), which substantially restored cellular proliferation (Figure 3D). These results underscore the critical role of GLUT1-mediated glucose utilization in sustaining redox homeostasis for the proliferation and survival of HNSCC.

**Figure 3.**
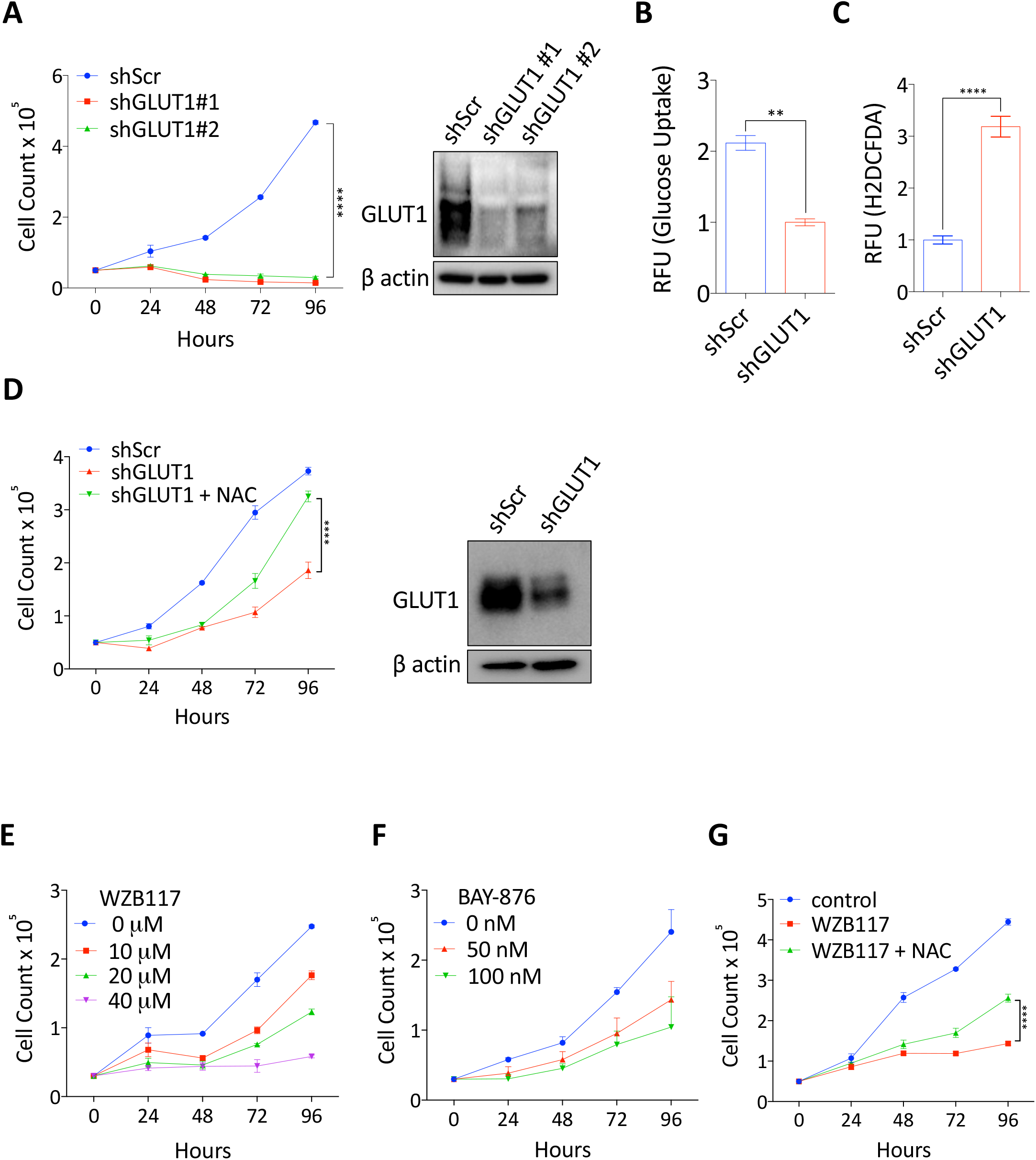
GLUT1 inhibition induces cytotoxic oxidative stress in HNSCC cells. (A) *in vitro* proliferation and immunoblot analyses of control shScramble (shScr) and shRNA-mediated knockdown of GLUT1 (shGLUT1) in HNSCC, FaDu cells; (n=4) Two-way ANOVA. (B) Relative glucose uptake and (C) intracellular ROS measurement in control shScr and shGLUT1 FaDu cells (n=3), *t*-test. (D) *in vitro* proliferation of control shScr, shGLUT1 and shGLUT1 + N-acetylcysteine-treated (NAC, 5 mM) HNSCC SCC61 cells; and immunoblot validation shRNA-mediated knockdown of GLUT1 in SCC61 cells (n=4; Two-way ANOVA. (E – F) *in vitro* proliferation of FaDu cells treated with GLUT1 inhibitor, WZB117 (0 – 40 μM) (E) or GLUT1 inhibitor, BAY876 (0 – 100 nM) (F). (G) *in vitro* proliferation of SCC61 cells treated with WZB117 (40 μM) and NAC (5 mM) (n=4; Two-way ANOVA). \*\*\*\**P*<0.0001, \*\**P*<0.01.

Furthermore, to validate glucose reliance of HNSCC cells, we pharmacologically inhibited GLUT1-mediated glucose influx using well-characterized irreversible GLUT1 inhibitors WZB117 and BAY-876 (30,31). Treatment with these inhibitors resulted in a dose-dependent decrease in HNSCC cell proliferation *in vitro* (Figure 3E – 3F), which could be significantly rescued by NAC supplementation (Figure 3G). These findings provide evidence that GLUT1-mediated glucose utilization is essential for combating oxidative stress in HNSCCs (9).

### Combination of GLUT1 Inhibition and Ascorbate Suppresses HNSCC Tumor Growth

Given the potential vulnerability to oxidative stress upon GLUT1 inhibition, we reasoned that combining GLUT1 inhibition with a strategy that induces cellular oxidative stress could elicit superior cytotoxic activities in HNSCC. Pharmacologic concentrations of vitamin C have been shown to function as a prooxidant and exert cytotoxic effects on various cancer types, including SCC (32,33). Indeed, treatment of HNSCCs with a combination of GLUT1 inhibitor, WZB117 (20 μM) and vitamin C (0.5 mM) resulted in a dramatic reduction of cellular proliferation as compared to either monotherapy treatment, thereby demonstrating synergy (Figure 4A). The combination of WZB117 and vitamin C treatment elicited the highest levels of intracellular H_2_O_2_, thereby implicating ROS and increased oxidative stress in decreasing cellular proliferation (Figure 4B).

**Figure 4.**
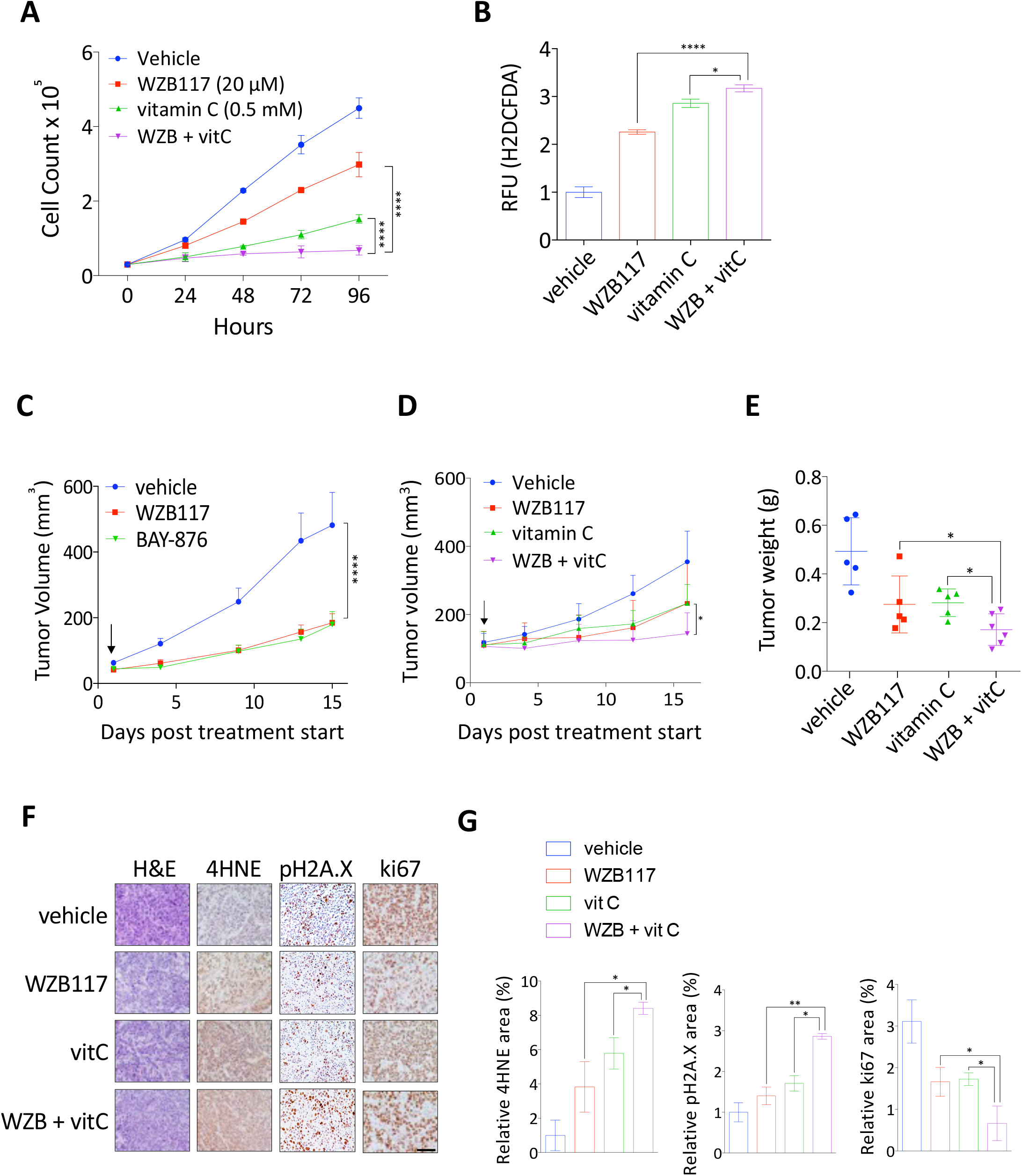
Combination of GLUT1 inhibition and ascorbate suppresses HNSCC tumor growth. (A) *in vitro* proliferation of FaDu cells treated with WZB117 (20 μM), vitamin C (0.5 mM) or combination of WZB117 and vitamin C (n=4; Two-way ANOVA). (B) Intracellular ROS measurement in control, WZB117 (20 μM), vitamin C (0.5 mM), and combination of WZB117 and vitamin C-treated FaDu cells (n=3; One-way ANOVA). (C) *in vivo* tumor growth of FaDu xenograft tumors treated with vehicle PBS/DMSO, WZB117 (20 mg/Kg i.p. daily) or BAY876 (4.5 mg/Kg corn oil/DMSO oral gavage daily) (n=6; Two-way ANOVA). (D) *in vivo* tumor growth, (E) tumor weights, and (F) IHC analysis of 4HNE, Ki67 and pH2AX in SCC61 xenograft tumors treated with vehicle PBS/DMSO (n=6), WZB117 (20 mg/Kg, i.p. daily) (n=5), vitamin C (4 g/Kg, i.p. daily) (n=5) and combination of WZB117 and vitamin C (n=6). Two-way ANOVA. Scale bar = 1mm. (G) IHC analysis quantification of % area of 4HNE (lipid peroxidation marker), pH2A.X (DNA damage marker) and ki67 (nuclear/cell proliferation marker) in xenografted tumors. 4 to 6 images of each tumor were captured and analyzed for quantification. One way ANOVA. \*\*\*\**P*<0.0001,\*\**P*<0.01, \**P*<0.05.

To further evaluate the suppressive effects of redox perturbation by either GLUT1 inhibition or the combination of prooxidants on HNSCC tumor growth *in vivo*, we treated SCID mice bearing human HNSCC FaDu xenograft tumors with GLUT1 inhibitors WZB117 (10 mg/kg/day) and BAY-876 (4.5 mg/kg/day) for 2 weeks. Both WZB117 and BAY-876 significantly reduced HNSCC xenograft tumor growth compared to the vehicle-treated mice (Figure 4C). Furthermore, consistent with our findings *in vitro,* treatment of mice with a combination of WZB117 and vitamin C (4 g/kg/day) resulted in significantly decreased HNSCC xenograft tumor growth and tumor weights compared to either of the monotherapy groups (Figure 4D – 4E). Immunohistochemical analysis of oxidative stress markers, 4HNE (lipid peroxidation) and pH2A.X (oxidative DNA damage), indicate significantly higher intratumoral oxidative stress, in HNSCC xenograft tumors treated with combination of WZB117 and vitamin C as compared to monotherapy tumors (Figure 4F – 4G). These results highlight the efficacy of combinatorial redox perturbation as a viable treatment modality for HNSCC.

### Auranofin induces cytotoxicity as a prooxidant in HNSCC

To explore potential drug candidates that exploit redox vulnerabilities in HNSCC, we utilized the Drug Repurposing Hub (Broad Institute, https://clue.io/repurposing) to search for FDA-approved drugs targeting key antioxidant enzymes and functioning as prooxidants for cancer therapy. Amongst the options, we selected auranofin, an FDA-approved drug primarily used for the treatment of rheumatoid arthritis (34). Auranofin has been shown to inhibit thioredoxin reductase 1 (TrxR), an antioxidant enzyme that acts on electron cycling in the NADPH/GSH electron coupling system. We hypothesized that inhibition of TrxR with auranofin could potently disrupt the cellular redox balance and precipitate oxidative stress in HNSCC cells (27, 28). Indeed, treatment of HNSCCs with auranofin resulted in decreased cellular proliferation and increased oxidative stress (Figure 5A – 5B). Consistent with ascorbate, auranofin elicited robust anti-proliferative activities when combined with GLUT1 inhibitor, WZB117 (Figure 5C).

**Figure 5.**
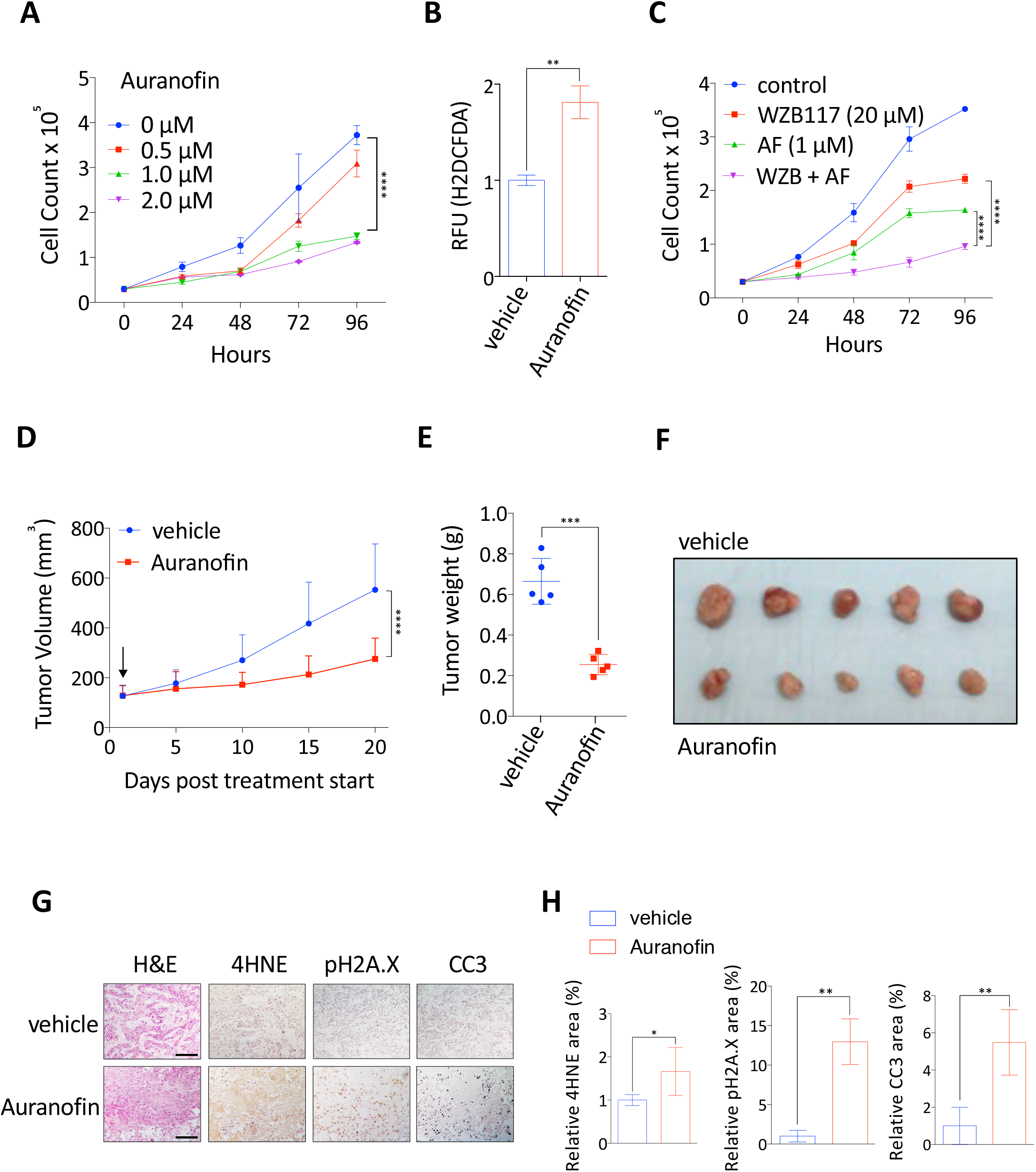
Auranofin induces cytotoxicity as a prooxidant in HNSCC. (A) *in vitro* proliferation of SCC61 cells treated with 0 – 2 μM Auranofin. (B) Intracellular ROS measurement in vehicle or Auranofin (1 μM)-treated SCC61 cells (n=3; *t*-test). (C) *in vitro* proliferation of SCC61 cells treated with vehicle, WZB117 (20 μM), Auranofin (2 μM) or combination of WZB117 and Auranofin (n=4; Two-way ANOVA). (D) *in vivo* tumor growth (two-way ANOVA), (E) tumor weights (*t*-test), (F) individual tumors and (G) IHC analysis of GLUT1, pH2AX, 4HNE, and CC3 in SCC61 xenograft tumors treated with vehicle PBS/DMSO (n=6) or Auranofin (10 mg/Kg, i.p. daily) (n=6) (H) IHC analysis quantification of % area of 4HNE (lipid peroxidation marker), pH2A.X (DNA damage marker) and cleaved caspase-3 (CC3; apoptotic cell marker) in xenografted tumors. 4 to 6 images of each tumor were captured and analyzed for quantification. Two-tailed *t*-test. \*\*\*\**P*<0.0001, \*\**P*<0.01, \**P*<0.05. Scale bars = 1 mm.

Next, we evaluated the preclinical efficacy of auranofin in HNSCC xenograft tumors. Daily intraperitoneal injection of auranofin (10 mg/kg/day) significantly inhibited tumor growth compared to the vehicle control group (Figure 5D – 5E). Immunohistochemical analysis indicated that auranofin treatment led to an increase in oxidative DNA damage, lipid peroxidation, and apoptotic cell death as indicated by pH2A.X, 4HNE, and cleaved-caspase 3 (CC3) staining, respectively, in HNSCC tumors (Figure 5F – 5G). These findings collectively highlight the potential of auranofin as a prooxidant that inhibits HNSCC tumor growth, positioning it as a promising candidate for HNSCC therapy, either as a standalone treatment or in combination with other drugs such as GLUT1 inhibitors.

### Insulin signaling promotes glucose-mediated antioxidative capacities in HNSCC

As glucose uptake and metabolism depend on insulin signaling, we aimed to investigate the impact of insulin signaling on tumor growth by modulating glucose-mediated antioxidative activities in HNSCC. To assess this, we first exposed human HNSCC cells to exogeneous insulin in serum-free conditions, which promoted proliferation and AKT activation, highlighting, perhaps not surprisingly, that insulin acts as a significant growth factor in HNSCC cells (Figure 6A – 6B). Intriguingly, insulin treatment resulted in a notable reduction in intracellular H_2_O_2_ levels likely attributed to elevated glucose influx in the context of prior results (Figure 6C – 6D). To further validate the role of insulin signaling in promoting HNSCC antioxidant capacity, we performed shRNA-mediated knockdown of the insulin receptor (IR/INSR). Insulin receptor knockdown led to decreased glucose uptake, increased oxidative stress, and inhibited proliferation of INSR knockdown cells was partially restored by NAC treatment, suggesting that insulin signaling plays critical roles for glucose utilization and redox homeostasis in HNSCC. However, it is important to note that pro-proliferative role of insulin signaling is not limited solely to its pro-antioxidant effects given that antioxidant supplementation only partially rescued growth upon INSR ablation.

**Figure 6.**
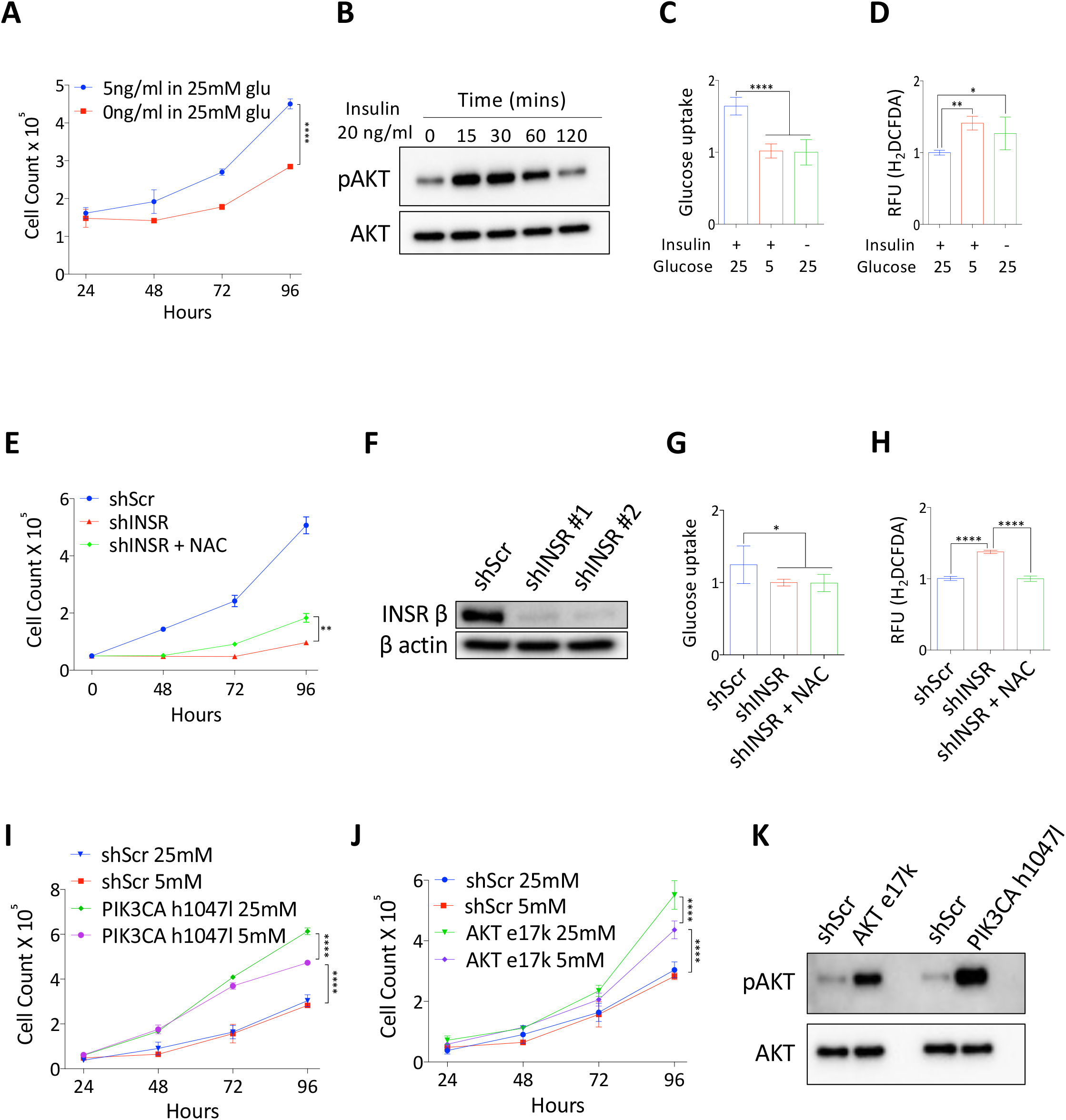
Insulin signaling promotes glucose-mediated antioxidative capacities in HNSCC. (A) *in vitro* proliferation of HNSCC FaDu cells in 25 mM glucose media ± 5 ng/mLl insulin (n=4, Two-way ANOVA). (B) Immunoblot analysis of pAKT and AKT expression in FaDu cells treated with 20 ng/mL insulin over a time period of 0 – 120 minutes. (C) Relative glucose uptake and (D) intracellular ROS measurement in FaDu cells treated with 20 ng/mL insulin in 25 mM or 5 mM glucose conditions (n=3; One-way ANOVA). (E) *in vitro* proliferation of control shScr, shINSR, or shINSR + NAC (5 mM) FaDu cells (n=4, Two-way ANOVA). (F) immunoblot analyses of control shScr and shINSR FaDu cells. (G) Relative glucose uptake and (H) intracellular ROS measurement in control shScr and shINSR FaDu cells (n=3; *t*-test). (I) *in vitro* proliferation of control shScr, oncogenic mutant AKT e17k and PIK3CA h1047l expressing FaDu cells grown in 5mM or 25mM glucose media (n=4, Two-way ANOVA). (J) immunoblot analysis of pAKT and AKT expression in shScr and mutant (AKT e17k and PIK3CA h1047l) FaDu cells. \*\*\*\**P*<0.0001, \*\**P*<0.01, \**P*<0.05.

Additionally, we introduced constitutively active mutants of the insulin signaling pathway: PIKC3A-H1047L and AKT-E17K. HNSCC cells expressing these mutants exhibited significantly enhanced proliferation (Figure 6I – 6J). Notably, lower glucose conditions (5 mM) restrained the proliferative capacity of these cells, presumably due to decreased glucose uptake and increased cellular oxidative stress (Figure 6I – 6J). Importantly, it is worth noting that lowering glucose to 5 mM had no effect on parental HNSCC cells, indicating that insulin signaling promotes cellular proliferation, in part, by augmenting glucose-mediated antioxidative activities in HNSCCs.

### Dysregulated insulin signaling reduces HNSCC tumor growth

To gain broader insight into how insulin signaling impacts the glucose utilization and tumor growth of HNSCCs *in vivo*, we employed a streptozotocin (STZ)-induced type I diabetes mouse model. This model involves the administration of STZ, which leads to the depletion of pancreatic beta cells and a subsequent decrease in insulin production (Figure 7A – 7B), resulting in the induction of type I diabetic phenotypes in mice (35). Intraperitoneal Glucose Tolerance Tests (IPGTT) confirmed severe disruption of glucose homeostasis, as evidenced by sustained high levels of blood glucose (> 250 mg/dL) in STZ-treated mice (Figure 7C). Remarkbly, HNSCC tumor growth in STZ-treated mice was significantly reduced compared to the control group (Figure 7D). This decrease in tumor growth was accompanied by an increase in intratumoral oxidative stress (Figure 7E – 7F). These findings strongly support the crucial role of insulin signaling in sustaining augmented glucose utilization for HNSCC tumor growth.

**Figure 7.**
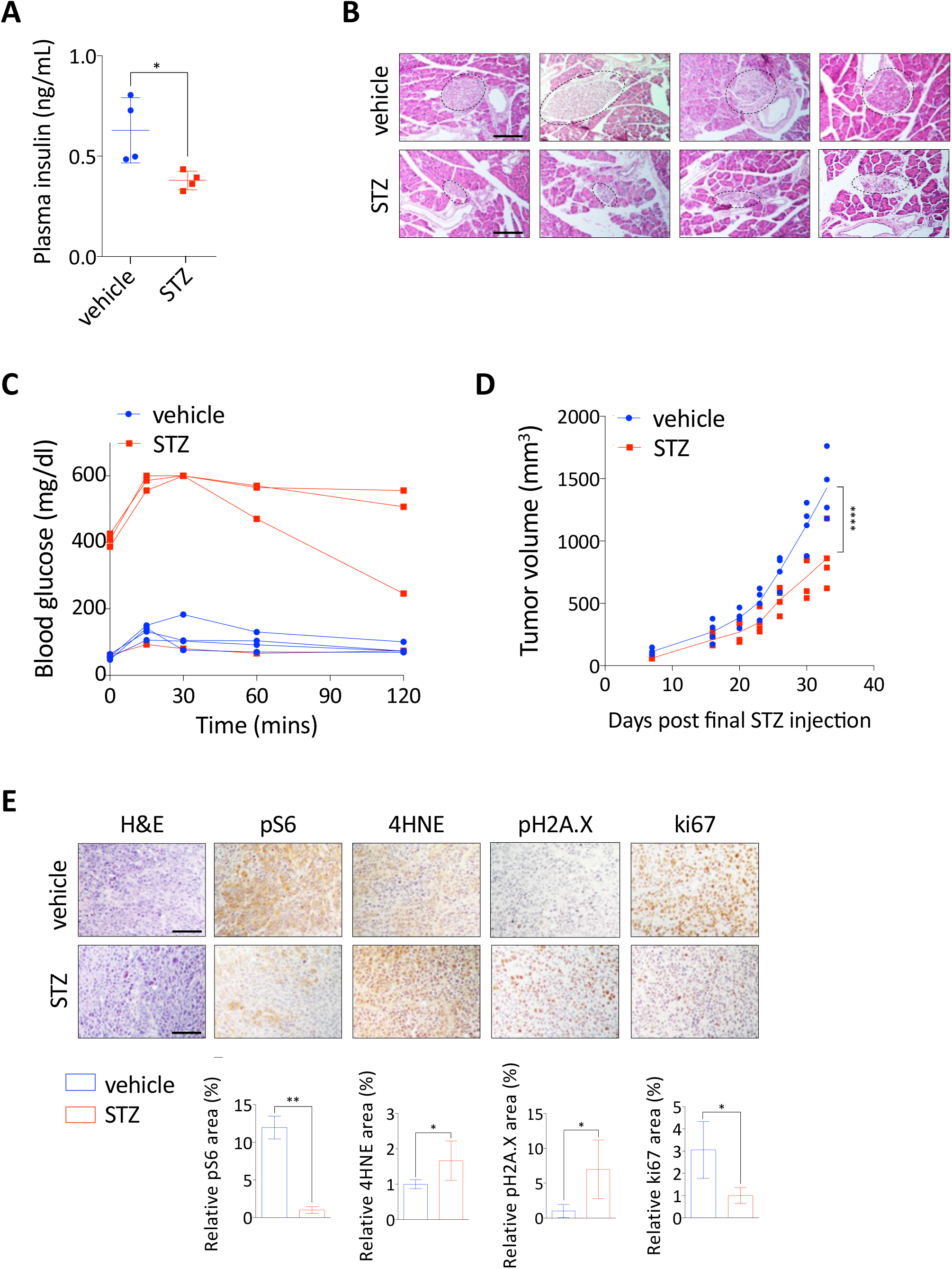
Dysregulated insulin signaling reduces HNSCC tumor growth. (A) Plasma insulin concentration of vehicle- and STZ (streptozotocin)-treated mice (40 mg/Kg i.p. for 5 consecutive days) (n=4; *t*-test) (B) representative H&E staining of pancreatic tissues collected from vehicle and STZ-treated mice. Dashed lines denote individual pancreatic islets. Scale bars = 1mm (C) intraperitoneal glucose tolerance test (IPGTT) of vehicle- and STZ-treated mice (n=4). (D) *in vivo* tumor growth of FaDu xenograft tumors in vehicle- and STZ-treated mice (n=4; Two-way ANOVA) (E) H&E and IHC analyses of pH2AX, 4HNE, and ki67 in FaDu xenograft tumors collected from vehicle and STZ-treated mice. (F) IHC analysis quantification of % area of 4HNE (lipid peroxidation marker), pH2A.X (DNA damage marker) and ki67 (nuclear/cell proliferation marker) in xenografted tumors. 4 to 6 images of each tumor were captured and analyzed for quantification. Two-tailed *t*-test, Scale bars = 1 mm. \*\*\*\**P*<0.0001, \*\**P*<0.01, \**P*<0.05.

## DISCUSSION

HNSCC poses a significant challenge due to limited treatment options and frequent late-stage diagnoses, leading to high mortality rates. HNSCCs arise from a diversity of anatomical contexts and thereby vary in their molecular characteristics and mutational signatures, thereby resulting in both inter- and intra-tumoral cellular heterogeneity, which invariably affect therapeutic responses, outcomes and prognosis (36,37). The added complexity of the HNSCC metabolic landscape has likely impeded the development of targeted therapeutics. Despite traditional therapies such as surgery, chemotherapy, and radiotherapy, HNSCC patients often experience high rates of relapse with locoregional disease and widespread metastases emphasizing the urgent need to identify key oncogenic events that are necessary for continued progression and promote HNSCC malignancy. In this study, we establish GLUT1-mediated glucose influx and associated redox metabolism, as a functional driver of malignant progression in HNSCC, and therefore as a critical metabolic feature that can be targeted for diagnosis and therapy. By connecting insulin signaling and glucose metabolism to the regulation of redox homeostasis, which we demonstrate as essential for SCC survival, we define a unifying metabolic phenotype amongst squamous cancers that can be exploited to achieve improved patient care.

Our initial analysis of a diverse set of HNSCC patient tumor samples from various anatomical origins (pharyngeal, laryngeal and other oral cancers) in TCGA data revealed that HNSCCs exhibit remarkable overexpression of the glucose transporter GLUT1. Using the mouse model of human HNSCC induced by 4NQO1, which recapitulates the pro-oncogenic effects of cigarette smoke or alcohol by promoting oxidative DNA and cellular damage within squamous epithelial cells of the tongue and esophagus, we observed dramatic GLUT1 upregulation in pre-malignant squamous lesions (Figure 2B). This suggests that GLUT1-mediated glucose influx and utilization play a crucial role in facilitating squamous malignant progression. These data support our previous discoveries demonstrating that glucose influx into squamous cells is utilized to mount a critical antioxidative response, simultaneously fueling and supporting uncontrolled cancer progression (9). To target this glucose influx in HNSCC, we employed multiple approaches, including GLUT1 knockout in HNSCC cells of origin, GLUT1 knockdown, and pharmacological inhibition of GLUT1 both *in vitro* and *in vivo*. All approaches led to a decrease in GLUT1-mediated glucose utilization for antioxidative capacities as indicated by elevated ROS, severely impairing cancer growth. This strongly supports targeted disruption of redox homeostasis as a viable strategy for HNSCC therapy. Notably, of the combination of prooxidant high-dose vitamin C and auranofin exhibited synergistic cytotoxicity when used in conjunction with GLUT1 inhibition.

Considering the crucial impact of insulin signaling on central carbon metabolism, we sought to understand the overall impact of dysregulated insulin signaling on HNSCC tumor progression. Clinical and epidemiological studies have implicated diabetes with an increased incidence of HNSCC and poorer prognosis for HNSCC patients (38–40). Yet, how insulin signaling regulates glucose-mediated redox metabolism remains largely unexplored. Building upon our recent findings demonstrating that a ketogenic (low carbohydrate, high fat) diet was able to significantly reduce lung squamous cancer survival and progression via restriction of glucose availability and indirect suppression of insulin signaling (9), our current study demonstrates that HNSCC cells robustly respond to insulin by increasing cellular proliferation, glucose uptake, and antioxidative capacities. Knockdown of the insulin receptor resulted in severe cytotoxicity in HNSCC cells, which was partially rescued by the antioxidant NAC. Furthermore, HNSCC tumor growth was significantly impaired in a hypoinsulinemic condition induced by STZ, which was associated with increased intratumoral oxidative stress. These findings suggest that insulin plays a critical role in the proliferation and tumor growth of HNSCC in part by augmenting glucose-fueled antioxidative capacities.

Recent advances in therapeutic techniques, such as targeted therapies and immunotherapy, offer exciting alternatives to drastically decrease the challenges of toxicity and hazardous side effects of traditional therapies. Clinical trials investigating immunotherapy and targeted therapy have shown some therapeutic promise (7,41–44). In this regard, several molecular targets, including epidermal growth factor receptor (EGFR), vascular endothelial growth factor (VEGF), and phosphatidylinositol-3-kinase (PI3K), have been implicated in the treatment of HNSCC (45–47). However, overall clinical outcomes have not yet indicated conclusive therapeutic benefits. Our data strongly support the rationale for targeting glucose-induced redox homeostasis in HNSCC and other SCCs. In particular, we propose the use of auranofin to achieve these ends, although further validation of auranofin as a candidate therapeutic for squamous cancer patients is needed. Auranofin is currently being tested in ongoing clinical trials for multiple cancer types (ovarian cancer NCT03456700, chronic lymphocytic leukemia NCT03456700 and lung cancer NCT03456700) supporting its potential translatability for HNSCC patients.

In conclusion, this study demonstrates promising preclinical efficacy using multifaceted approaches to induce cytotoxic oxidative stress and impede the tumorigenic progression of HNSCC. These findings highlight the potential of targeting redox vulnerabilities as an effective therapeutic option for HNSCC patients, warranting further investigation of the clinical implications of prooxidant therapy.

## MATERIALS AND METHODS

### Mice

C57BL/6 and NOD/SCID mice were obtained from Jackson Laboratory. All experimental procedures utilizing mice were approved by The University of Texas at Dallas Institutional Animal Care and Use Committee

### 4-nitroquinoline 1-oxide (4NQO)-induced head & neck and esophageal cancer model

4NQO was dissolved in propylene glycol 10 mg/mL and diluted into the drinking water to 100 ug/mL (27). Mice were treated with 4NQO for a maximum treatment time of 8 weeks and then observed for up to an additional 20-week post treatment. Briefly, after 8 weeks, 4QNO was removed from the drinking water and being maintained with normal drinking water for additional 4 weeks (total 12 weeks after initial exposure to 4NQO) Mice were then injected intraperitoneally (i.p.) with tamoxifen (100 mg/Kg once per day for 5 consecutive days) to delete GLUT1 gene in the Krt5^+^ basal progenitor cells. dTomato, included as a fluorescent reporter of Cre activity in our model system, was activated specifically in GLUT1-expressing basal epithelial cells upon treatment of mice with tamoxifen. Control mice were given normal water with a vehicle of propylene glycol for the entire treatment period. All mice were given water + 4NQO or water + vehicle *ad libitum*. During treatment period animals were monitored daily and weighed weekly. The oral mucosa of each mouse was examined under inhalation anesthesia weekly to check for oral tumors. Food intake was closely monitored for animals developing esophageal tumors. Mice were euthanized immediately when they showed signs of stress, weight loss, or discomfort. To provide a comprehensive temporal analysis of cancer progression and severity, mice were euthanized, and tongues and esophagi were collected for analysis.

### Basal-epithelial-cell-specific GLUT1 knockout

Keratin 5 (Krt5) is an intermediate filament protein specifically expressed in the basal layer of stratified epithelial cells from which premalignant dysplastic squamous lesions arise. To generate a conditional deletion of GLUT1 in the basal squamous epithelium KRT5:Cre mice were crossed with GLUT1 floxed mice. We also introduced a heterozygous deletion of the tumor suppressor p53, which mimics loss of heterozygosity common in HNSCC. In our model: *Krt5:CreER; LSL-dTomato; P53KO+/-; GLUT1 ^fl/fl^* mice express tamoxifen-inducible Cre recombinase under the direction of the *KRT5* promoter. Cre activity was assessed with a fluorescent tomato reporter dTomato. After the administration of tamoxifen (100 mg/Kg once per day for 5 consecutive days), the gene encoding GLUT1 was knocked out in the basal, KRT5+ squamous epithelium.

### Cell lines

SCC61 HNSCC cell line was provided by Jeffrey Myers (MD Anderson Cancer Center). FaDu HNSCC cell line was purchased at ATCC (HTB-43). Cells were cultured in high glucose DMEM (Sigma) supplemented with 10% fetal bovine serum (Sigma), 1% penicillin/streptomycin (Sigma) and 1% non-essential amino acids (Sigma) (unless otherwise stated) at 37°C in a humidified 5% CO_2_ environment. All cell lines were mycoplasma tested with e-Myco Kit (Boca Scientific).

### Prooxidant vitamin C experiments

To pharmacologically inhibit GLUT1 we used the well-characterized irreversible GLUT1 inhibitors WZB117 and BAY-876 to target GLUT1-mediated glucose influx and its related function in redox homeostasis (30,31,48,49). The specificity of these GLUT1 inhibitors has been previously well validated and demonstrated in our previous studies. We combined these GLUT1 inhibitors with prooxidant high-dose vitamin C to potently disable cellular oxidative defense machinery. Various cell lines were treated with WZB117 (10 – 50 μM) or BAY-876 (10 – 100 nM) and with vitamin C (0 - 1 mM). Following treatment glucose influx was evaluated using glucose uptake assays. Cellular oxidative stress will be examined by measuring intracellular H_2_O_2_ (H_2_DCFDA staining).

### In vivo tumor xenograft experiments

5X10^6 cells suspended in 50% matrigel (Corning Life Sciences) and 50% Hank’s Balanced Salt Solution (HBSS, Sigma) were subcutaneously implanted into the flank of NOD/SCID mice (Jackson Laboratory) between 4 and 6 weeks old. Cisplatin (Sigma) was administered intraperitoneally (i.p.) at 2 mg/Kg in PBS (Sigma) twice weekly. WZB117 (Calbiochem) 10 mg/Kg was administered i.p. once daily. Tumor volume was measured at indicated times using electronic calipers and estimated by the modified ellipsoid formula: tumor volume = (length x width^2^)/2.

### GLUT1 inhibition *in vivo*

Mice were divided into four groups (6 - 12 mice each) and received i) vehicle control, ii) WZB117 (10 – 20 mg/Kg/day, i.p.) or BAY-876 (3 – 5 mg/Kg/day, oral gavage) only, iii) vitamin C (4 g/Kg/day, i.p.) only, and combination of WZB117/BAY876 and vitamin C. Subsequently to evaluate physiological effects of our proposed treatments, xenografts were established in SCID mice.

### Streptozotocin (STZ) diabetes mouse model

Six-week-old mice were individually marked, weighed, and their baseline blood glucose levels determined using a *OneTouch Ultra* blood glucose monitor. Next, mice received daily IP injections of 40 mg STZ/Kg body weight (for NSG & NOD-scid males) for five consecutive days; age-matched controls received buffer–only injections. STZ was dissolved in sodium citrate buffer. Food and water was available *ad libitum*. The mice were observed daily and blood glucose measurements were taken one week after the final injection. Mice with blood glucose levels exceeding 250 mg/dL were considered diabetic. HNSCC cell line xenografts were then established in diabetic and control mice. Tumor measurements were made for up to 3 weeks at which point tumors were excised for histopathologic, protein and RNA analyses. Prior to final tumor measurement, mice were evaluated to see if they are still diabetic. Plasma will also be collected for Insulin ELISA analysis.

### Blood glucose and insulin measurement

Blood collected from the tail of mice fasted for six hours prior was utilized to measure blood glucose via glucometer (ONETOUCH Ultra2). Up to 200 mL of blood was collected from the tail into EDTA coated microfuge tube and centrifuged to isolate plasma for insulin measurement. Insulin levels were determined via Mouse Insulin ELISA Assay Kit (Crystal Chem) according to the manufacturer’s instruction

### Immunoblotting

Cells were lysed by RIPA lysis buffer supplemented with cOmplete Protease Inhibitor (Roche) and subsequent 20% amplitude sonication for 5s, and lysates were cleared by 14,000 rpm centrifugation at 4°C for 15 min. Equivalent lysates were separated SDS PAGE and transferred onto polyvinylidene difluoridemembranes (FisherScientific). Membranes following blocking in 5% non-fat dry milk dissolved in TBST for 30 min were incubated in primary antibody diluted in 5% BSA overnight. Horseradish-peroxidase conjugated secondary antibodies diluted 1:5000 in 5% non-fat milk were used and visualized with SuperSignal West Pico or Femto substrate kits (ThermoFisher). The following commercial primary antibodies supplemented with 0.02% sodium azide were used for immunoblot analysis: p63 (1:1,000; Biocare Medical CM163A), GLUT1 (1:1,000; Alpha Diagnostics GT11-A), Tyr1361-pINSR (1:1,000; ThermoFisher Scientific PA5-38283), INSR (1:1,000; ThermoFisher Scientifc AHR0271), Ser473-pAKT (1:1000; Cell Signaling Technology #4058), pAKT (1:1,000; Cell Signaling Technology #9272), Ser235/236-pS6 (1:1,000; Cell Signaling Technology #4858), S6 Ribosomal Protein (1:1,000; Cell Signaling Technology #2217), Thr37/46-p4EBP1 (1:1,000; Cell Signaling Technology #2855), Cleaved Caspase-3 (1:1,000; Cell Signaling Technology #9664), and α-actin (1:5,000; Sigma A5441).

### Immunohistochemistry and immunofluorescence

Xenograft mice were perfused with 10mM EDTA in PBS followed by 4%PFA. Xenograft tumors were extracted and fixed in 4% PFA for 12 hours and were then embedded in paraffin. Tissue blocks were then sectioned (5μm) and subjected to heat-mediated antigen retrieval (citrate buffer, pH 6.0). Goat serum (Sigma) or donkey serum (Sigma) was used to block for 1 hour, and primary antibodies diluted were applied at 4C overnight. Vectastain ABC (Vector Labs) with DAB substrate (Vector Labs) was used to optimize staining according to the manufacture’s protocol. The following primary antibodies were used: p63 (1:200; Biocare Medical; CM163A), p63 (1:100; R&D Systems AF-1916), GLUT1 (1:250; Alpha Diagnostic GT11-A), Ki67 (1:500; Cell Signaling Technology #12202), Cleaved Caspase-3 (1:200; Cell Signaling Technology #9664), Ser473-pAKT (1:500; Cell signaling Technology #4058), Ser235/236-pS6 (1:200; Cell Signaling Technology #4858), Ser139-pHistone H2A.X (1:1,000; Cell Signaling Technology #9718), 4-Hydroxynonenal (1:500;Abcamab46545), CK5 (1:200;Abcamab52635). Images were taken via Nikon Eclipse Ni-U microscope with NIS Elements imaging software (Nikon) and quantified using ImageJ (NIH).

### *In vitro* ROS measurement

ROS levels were detected via H_2_DCFDA (Cayman) according to the manufacturer’s protocol. Following seeding on 96-well plates cells were stained with H2DCFDA for 1 hour. Reduced and oxidized probes were measured respectively at 590 nm and 535 nm. Emission at 535 nm was measured using a fluorescent confocal microscope (Nikon Eclipse Ni-U).

### In vitro metabolic analysis

Glucose uptake was measured using the Glucose Uptake Cell-Based Assay Kit (Cayman) according to the manufacturer’s protocol. Following incubation with fluorescent glucose analog2-NBDG in glucose-free DMEM (GIBCO) at 37°C for at least 1 hour and cell preparation according to the manufacturer’s instruction, emission at 535 nm was measured using a fluorescent confocal microscope (Nikon Eclipse Ni-U).

### shRNA and vectors

Following pLKO.1 shRNA were used: shINSR#1 (Mission TRC shRNA, TRCN0000010523, Sigma), shINSR#2 (Mission TRC shRNA, TRCN0000000379, Sigma), shGLUT1 (Mission TRC shRNA, TRCN0000043583, Sigma), shIGFR1#1 (Mission shRNA, TRCN0000039677, Sigma), shIGFR1#2 (Mission TRC shRNA, TRCN0000000422, Sigma). For lentivirus production, HEK293T cells were transfected with viral packaging plasmids psPAX2 Addgene #12260) and pMD2.G (Addgene#12259), and pLKO.1 shRNA using Lipofectamine 3000 (Invitrogen). Cells were incubated with viral supernatant containing polybrene (8 mg/mL). pLKO.1-shScr was used as a control vector. Targeting sequences for all shRNAs are provided in Table S1. pLenti CMV GFP Blast (659–1) was a gift from Eric Campeau & Paul Kaufman (Addgene plasmid # 17445). pHAGE-PIK3CA-H1047L was a gift from Gordon Mills & Kenneth Scott (Addgene plasmid # 116499; http://n2t.net/addgene:116499; RRID:Addgene_116499). pHAGE-AKT1-E17K was a gift from Gordon Mills & Kenneth Scott (Addgene plasmid # 116103).

### Quantification and statistical analysis

Statistical analyses were performed using StatPlus, Version v5 (AnalystSoft Inc.) and GraphPad Prism 7.0 (GraphPad Software Inc.). All data are expressed as mean ± s.e.m or median ± the interquartile range unless noted otherwise. Two-tailed Student’s t test, one way ANOVA with multiple comparison post hoc test, Kruskal-Wallis nonparametric ANOVA, Chi-Square test and Mann-Whitney U test were used for hypothesis testing, and p values of 0.05 were considered significant. ****p < 0.0001, ***p < 0.001, **p < 0.01, *p < 0.05

## ACKNOWLEDGEMENTS

We thank all members of J. Kim and TH Kim labs for helpful discussion and critical reading of the manuscript. This work was supported by the Department of Defense Lung Cancer Research Program W81XWH-18-1-0439 to J.K. and T.H.K.

## AUTHOR CONTRIBUTIONS

Conception and Design, S.M., J.K and T.H.K.; Investigation and data acquisition, S.M.; Data analysis and interpretation, S.M., J.H.C., T.G.O., J.K and T.H.K; Technical and material support, P.K.S., J.K. and T.H.K; Writing and revision of manuscript, S.M., J.H.C., J.K., and T.H.K.

